# A graph-based approach to identify motor neuron synergies

**DOI:** 10.1101/2023.02.07.527433

**Authors:** Simon Avrillon, François Hug, Dario Farina

**Author notes:** **Correspondence:** Dario Farina, Imperial College London, London, UK, **e-mail:**.

## Abstract

Multiple studies have experimentally observed common fluctuations in the discharge rates of spinal motor neurons, which have been classically interpreted as generated by correlated synaptic inputs. However, so far it has not been possible to identify the number of inputs, nor their relative strength, received by each motor neuron. This information would reveal the distribution of inputs and dimensionality of the neural control of movement at the motor neuron level. Here, we propose a method that generates networks of correlation between motor neuron outputs to estimate the number of common inputs to motor neurons and their relative strengths. The method is based on force-directed graphs, the hierarchical clustering of motor neurons in the graphs, and the estimation of input strengths based on the graph structure. To evaluate the accuracy and robustness of the method, we simulated 100 motor neurons driven by a known number of inputs with fixed weights. The simulation results showed that 99.2 ± 0.6%, 94.3 ± 2.2 %, and 95.1 ± 2.7 % of the motor neurons were accurately assigned to the input source with the highest weight for simulations with 2, 3, and 4 inputs, respectively. Moreover, the normalised weigths (range 0 to 1) with which each input was transmitted to individual motor neurons were estimated with a root-mean-squared error of 0.11, 0.18, and 0.28 for simulations with 2, 3, and 4 inputs, respectively. These results were robust to errors introduced in the discharge times (as they may occur due to errors by decomposition algorithms), with up to 5% of missing spikes or false positives. We finally applied this method on various experimental datasets to demonstrate typical case scenario when studying the neural control of movement. Overall, these results show that the proposed graph-based method accurately describes the distribution of inputs across motor neurons.

**Authors summary:** An important characteristics for our understanding of the neural control of natural behaviors if the dimensionality in neural control signals to the musculoskeletal system. This dimensionality in turn depends on the number of synaptic inputs transmitted to the elementary units of this control, i.e., the spinal motor neurons, and on their correlation. We propose a graph-based approach applied to the discharge times of motor neurons to estimate the number of inputs and associated strength transmitted to each motor neuron. For this purpose, we first calculated the correlation between motor neuron smoothed discharge rates, assuming that correlated discharge rates result from the reception of a correlated inputs. Then, we derived networks/graphs in which each node represented a motor neuron and where the nodes were positioned close to each or further apart, depending on the level of correlated activities of the corresponding motor neurons. Using simulations of motor neuron behaviour, we showed that the spatial information embedded in the proposed graphs can be used to accurately estimate the number and the relative strengths of the inputs received by each motor neurons. This method allows to reconstruct the distribution of synaptic inputs to motor neurons from the observed motor neuron activity.

## Introduction

Even the simplest of human movements involve the coordinated activity of thousands of motor units to generate the desired muscle forces. To achieve multi-muscle coordination, numerous studies have suggested a robust and low-dimensional control of muscles through the combination of motor modules, or *muscle synergies* (1, 2, 3, 4). Muscle synergies are functional units that generate a motor output by imposing a specific activation pattern on a group of muscles (5, 6). Generally, their identification relies on the combination of electromyographic (EMG) recordings over numerous muscles with factorization algorithms that extract consistent patterns of activity across muscles (7). It has been hypothesized that each synergy would be associated to a single neural command, which would in turn decrease the computational load of movement control (4, 8, 9).

The concurrent developments of high-density EMG electrodes and algorithms that decompose EMG signals into individual motor unit spiking activity have opened new perspectives in the study of movement control (10). It is now possible to isolate the activity of tens of individual motor units from multiple muscles and to estimate their correlated activity (11, 12, 13, 14). Several previous works have shown that motor units from a single muscle (15, 16, 17, 18, 19) or multiple synergistic muscles (11, 12, 13, 15) exhibit common fluctuations of their discharge rates, likely due to the presence of correlated inputs. These observations are in agreement with the muscle synergy theory. However, it has also been observed that a pool of motor neurons innervating the same muscle do not always receive the same inputs (14, 20, 21, 22, 23, 24). Therefore, we recently proposed that the central nervous system may generate a movement by transmitting a few inputs to groups of motor units (25), rather than to entire pools innervating individual muscles. In this view, the study of human movement control should be done at the motor neuron level, using methods that can identify the structure of the common inputs transmitted to motor neurons, rather than at the global muscle level.

While the projection of multi-muscle EMG envelopes into low-dimensional spaces can be done using several factorization algorithms (7), the problem of the identification of the structure of inputs to motor neurons has not been extensively addressed. Here we propose a graph-based approach to identify the number and relative strength of the inputs received by each motor neuron (14, 26, 27). To test the accuracy of this approach, we used a realistic model of a population of 100 motor neurons driven by a known number of inputs with fixed weights for each motor neuron (28). The graph-based approach relied on the identification of significant correlations between individual motor neuron smoothed discharge rates to estimate whether they received common inputs (29). Then, an algorithm generated a force-directed graph by positioning motor neurons with correlated outputs close to each other, while motor neurons exhibiting low correlation in their output discharge times were positioned farther apart (30). We then identified the number of common inputs received by the full population of motor neurons by detecting the unconnected sections of the graph and we estimated the relative strength of the inputs for each motor neuron by measuring the distance that separated the motor neurons from the centroid of each unconnected cluster. We tested the robustness of this graph-based approach to altered versions of motor neuron spike trains that mimicked potential inaccuracies of the EMG decomposition approach, such as false identification of action potentials, missing spikes, or a limited number of identified motor neurons. Finally, we present applications of our approach on representative experimental data from various muscles.

## Methods

### A graph-based approach to identify motor neuron synergies

The general idea of this approach is to consider motor neurons as nodes connected with springs, such that they attract each other if they exhibit correlated activity (30). We propose here to take advantage of the spatial properties of this network of motor neurons to infer the number of inputs they receive and their relative strength for each motor neuron.

#### Overview of the simulation paradigm

We simulated 100 S-type motor neurons receiving correlated and independent synaptic inputs using a motor neuron model previously proposed (28). This model has two compartments for the soma and the dendritic tree, respectively, Na^+^, fast and slow K^+^ conductance that generate action potentials and afterhyperpolarization, with the addition of a L-type Ca^++^ channel into the dendritic compartment to consider the non-linearity between motor neuron inputs and outputs due to the activation of persistent inward currents (31). Geometric and electrical parameters of the 100 motor neurons were linearly interpolated from the range of values reported in Table 1 of (28). We defined the total input for each motor neuron as:

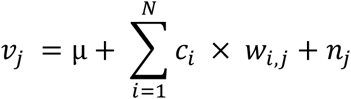

where μ is a constant offset that represents the mean of the synaptic inputs, N is the number of inputs, c_i_ is the temporal profile of the i^th^ input, w_i,j_ is the relative strength of the *i*^th^ input for the motor neuron *j*, and n_j_ is a synaptic input independent from the other motor neurons. We simulated *c* and *n* as Gaussian noises with equal variances. The inputs c had a bandwidth of 0-2.5Hz while the independent inputs had a bandwidth of 0-50Hz. Mean and variance of the synaptic current were tuned to generate physiologically realistic Poisson distributions of the interspike intervals (18).

We performed three simulations with N = 2, 3, and 4 inputs and a simulation where motor neurons only received independent inputs (Figure 1). For the simulation with N = 2, 40 motor neurons had either w_1,j_ or w_2,j_ set to 1, i.e., 20 motor neurons received only the first source of common input and 20 other motor neurons received only the second source of common input. For the 60 remaining motor neurons, w_1,j_ was linearly interpolated between 0.99 and 0.01, and w_2,j_ was equaled to 1 - w_1,j_. For the simulations with 3 or 4 correlated inputs, groups of 10 motor neurons received a single source of common inputs. The remaining motor neurons were grouped by 10 with the same set of w values. These values were randomly distributed across sources of correlated inputs to reach a total of 1. The duration of each simulation was set to 60 s.

**Figure 1:**
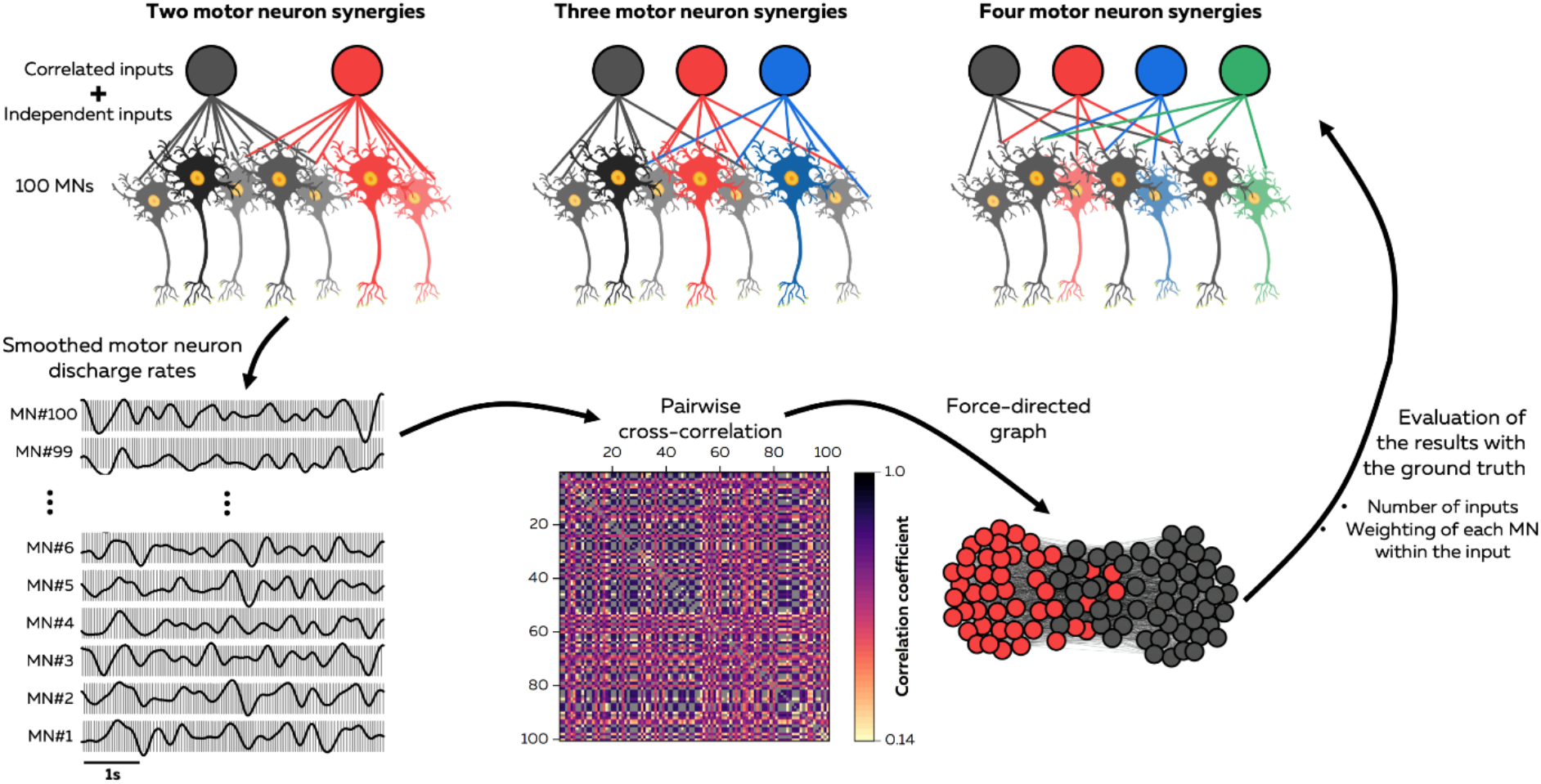
Overview of the simulation paradigm. We simulated the firing activity of 100 motor neurons receiving a known number of correlated inputs, i.e., 2, 3, and 4, with a relative strength consisting of a ratio between 0 and 1. Motor neurons firings were convoluted with a 400ms Hanning window to only consider the oscillations in the bandwidth of force fluctuations. Then, pair-wise cross-correlations were performed between the smoothed discharge rates, and correlation below a significance thereshold were discarded from the analysis. A force-directed graph was generated by applying the Fruchterman and Reingold algorithm (30) on the adjacency matrix of correlations. The graph has a number of nodes equal to the number of motor neurons, and each edge between motor neurons is a significant correlation. Finally, a hierarchical clustering algorithm (32) grouped motor neurons with a high level of correlation together, i.e., the red and grey nodes on this figure. We used both information on the clusters and the position of the nodes within the two-dimensional space to estimate the number of inputs and their weight for each motor neuron. Validation of our approach was done through comparison with the ground truth.

#### Correlation between motor neurons to identify common inputs

The output of each simulation was a matrix of 100 motor neuron spike trains organized row-wise and computed as binary vectors, where ‘ones’ were discharge instances. We computed the smoothed discharge rate of each motor neuron by convolving its spike train with a 400-ms Hanning window. The duration of the Hanning window was chosen to consider the fluctuations of motor neuron discharge rates effective for force modulation (16, 33). The smoothed discharge rates were then high-pass filtered with a cut-off frequency of 0.75 Hz to remove offsets and trends (Figure 1, (16)).

We assumed that pairs of motor neurons received common inputs when their smoothed discharge rates were significantly correlated (29). To identify these correlated activities, we performed cross-correlation on the smoothed discharge rates with a maximal time lag of 100 ms (16). We considered the statistical significance threshold as the 99^th^ percentile of the cross-correlation coefficient distribution generated with resampled versions of all motor neuron spike trains. To this end, we generated random spike trains for each motor neuron by bootstrapping the interspike intervals (random sampling with replacement). This random spike train had the same number of spikes and the same discharge rate (mean and standard deviation) as the original motor neuron spike train. We repeated this step four times per pair of motor neurons. At the end, all the correlation coefficients below the significance threshold were removed from the 100 by 100 matrix of correlation, which was used to generate the force-directed graph (Figure 1).

#### Force-directed graphs

We used graph theory to identify networks shaped by the correlated inputs received by all the motor neurons (14, 26, 27). Each graph had a set of 100 nodes, i.e., 100 motor neurons, and a set of edges, i.e., significant correlation between motor neuron activities, where each edge connected two nodes. We used the Fruchterman-Reingold algorithm implemented in the Python toolbox NetworkX (Release 3.0) to draw a graph using a force-directed placement of individual motor neurons into a two-dimensional space. Importantly, the position depended on the connectivity between individual motor neurons (30): each edge connecting two motor neurons was considered as a spring with attractive and repulsive forces depending on its length (30). In this way, motor neurons with correlated activity tended to be positioned closer to each other due to the attractive force of their edges, while the repulsive force avoided overlapping of motor neurons when the length of the edges was close to zero. The algorithm iteratively optimized node positions to minimize the total energy of the network. In practice, the graph converged to a spatial organization where groups of motor neurons with numerous edges were grouped together while groups of motor neurons with few edges were positioned evenly within the two-dimensional space.

#### Identification of the number of common inputs

After building the force-directed graphs, we applied a hierarchical consensus clustering procedure to group the motor neurons based on their correlated activity (14, 27). The aim of this approach was to identify groups of motor neurons densely connected to each other and loosely connected to the rest of the network (Figure 2). To this end, we used the multiresolution consensus clustering approach (32). First, we generated 1,000 partitions that covered the entire range of resolutions, i.e., from a partition where all the motor neurons belonged to the same cluster to a partition where the number of clusters was equal to the number of motor neurons. This approach ensures an approximately equal coverage of all scales of the network. Second, we applied consensus clustering on the entire set of partitions (34). This step is an iterative procedure that consists of: 1) estimating the probability for each pair of motor neurons to belong to the same cluster across the partitions, resulting in a consensus matrix; 2) identifying clusters within this consensus matrix with a graph-clustering algorithm that optimizes the modularity, i.e., the GenLouvain algorithm (32), to generate a new set of partitions; and 3) repeating 1) and 2) until the procedure converges toward a unique partition (34). This “consensus partition“ is considered as the most representative of all the partitions. It is also noteworthy that this algorithm only considers statistically significant consensus partitions. This means that the clusters of the ‘consensus partition’ cannot be identified in random networks generated by locally permutating the connections between motor neurons (32).

**Figure 2:**
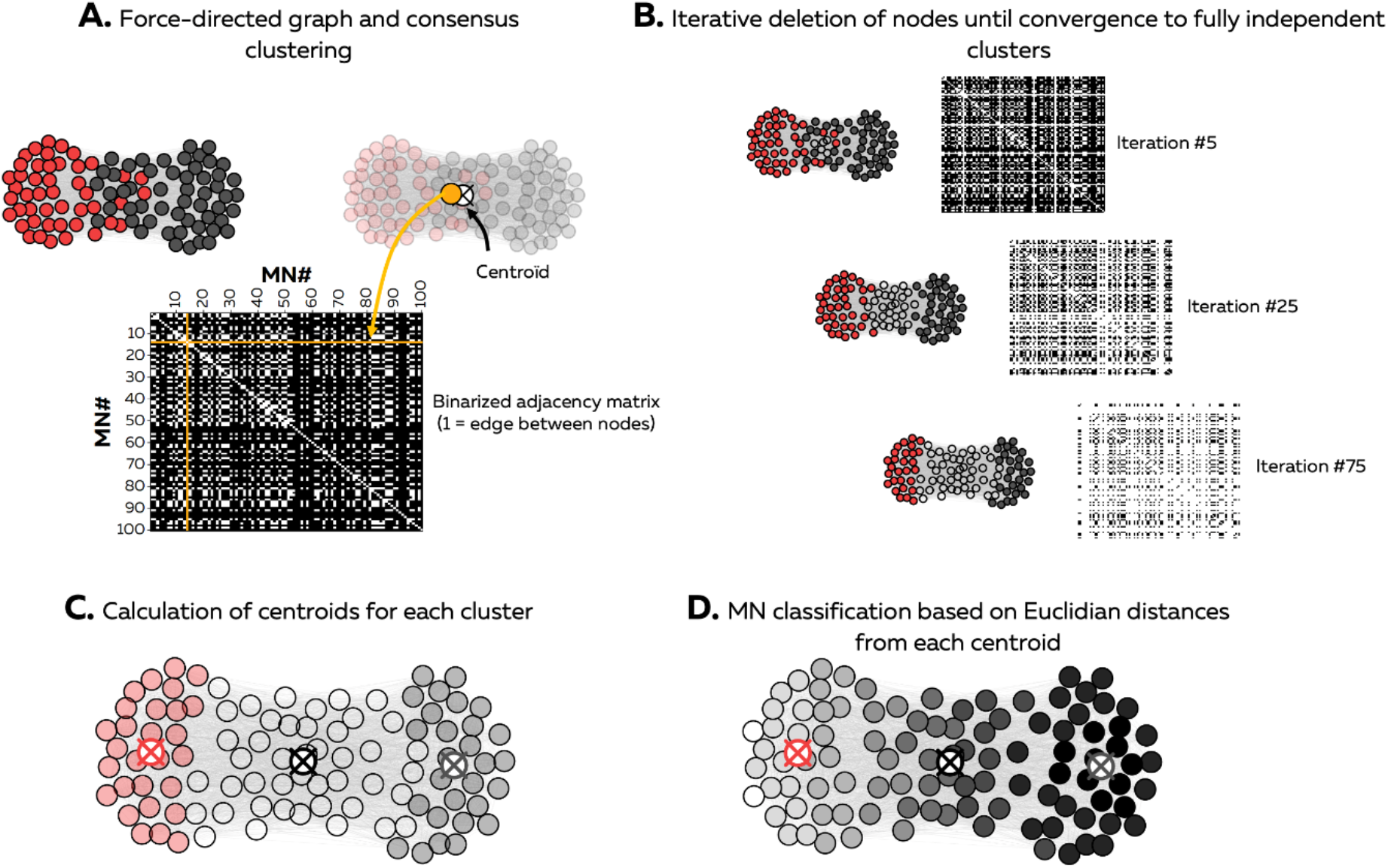
Estimation of the number of inputs and their relative strength from force-directed graphs. We used both information on the clustering of motor neurons and their position within the two dimensional space to estimate the number of inputs and their weighting. The iterative procedure started from the closest node from the centroid of the graph toward the farthest node. At each iteration, the algorithm selected all the motor neurons connected to the node (A). If these motor neurons belonged to different clusters, all the values of the vectors (yellow in A) were set to 0, and the motor neuron was discarded from the subsequent iterations (B). On the opposite, if these motor neurons belonged to the samer cluster, all the values from the vectors were kept (B). Thus, we only kept the groups of motor neurons that were fully independant to each other. We then calculated the centroid of each of these independant clusters (C), and calculated the distance of each motor neuron to these centroids (D). We considered the number of independant clusters as the number of inputs, and the distance between each motor neuron and the centroids as the relative strength of these inputs for each motor neuron.

To estimate the number of clusters in the graph (i.e., inputs), we identified fully independent groups of motor neurons, i.e., groups that did not share any connections or inputs with each other. While the algorithm used to draw the graph pushes these groups to the edges of the two-dimensional space, motor neurons that receive several correlated inputs tend to be positioned close to the center of this plane. Therefore, we performed an iterative procedure starting from the closest motor neuron from the centroid of the graph to the farthest. The algorithm deleted one motor neuron per iteration if it was connected to at least one motor neuron belonging to a different cluster. This procedure stopped when it converged to fully independent groups of motor neurons. Then, we estimated the weight for each motor neuron based on the Euclidean distance between the motor neuron and the centroid of each refined cluster. Indeed, as the position of each motor neuron within the two-dimensional space depends on the number of connections it shares with the others, the closer the motor neurons from each centroid, the greater the number of significant correlations they have with the motor neurons from this cluster. We performed this analysis for each cluster to obtain a vector of weights for each input. Finally, the Euclidean distances were normalized between 0 and 1 and subtracted from 1, given that the shorter the distance, the higher the number of significant correlations.

### Robustness and accuracy of the graph-based approach

We first verified the assumption that motor neurons with a significant correlation between their smoothed discharge rates received correlated inputs using the simulation with 2 inputs. To this end, we displayed the distributions of differences between the weights of input #1 for pairs with significant and non significant correlations. A difference equal to 1 means that one motor neuron receive 100% of input #1 while the other motor neuron receives 0% of input #1. On the opposite, a difference of 0 means that both motor neurons receive the same level of input #1. We repeated this analysis with correlations calculated over segments of 10s, 20s, 30s, 40s, 50s, and 60s to test the robustness of our approach across different durations of recordings.

To assess the robustness of the graph-based approach to identify the number of inputs and their relative strength for each motor neuron, we simulated perturbations of motor neuron spike trains that mimicked inaccuracies in the spike identification algorithms (EMG decomposition). First, we generated altered versions of each motor neuron spike trains by randomly removing or adding discharge instances. Specifically, we removed 1% to 9% of the total number of discharge instances or added false discharge instances equal to 1% to 9% of the initial number of discharge instances per motor neuron. We chose this range of values to get slightly lower sensitivity values that those generally observed after automatic EMG decomposition without manual edition (35). We then applied the graph-based approach as described above to identify the number of inputs and their relative weights for each motor neuron.

We also tested how the number of identified motor neurons impacted the estimation of the number of inputs and their relative weights. To this end, we randomly selected motor neurons out of the initial group of 100 motor neurons, starting from 10 motor neurons, and then increasing the size of the group by steps of 10, until reaching 90 motor neurons. We repeated this procedure 50 times per size of group.

To assess the accuracy of the graph-based approach, we first compared the number of identified inputs with the number of simulated inputs. Then, we computed the root-mean squared error (RMSE) between the estimated weights and the weights used in each simulation. Euclidean distances from the graph-based methods were normalized between 0 and 1 and subtracted from 1, given that the shorter the distance, the higher the number of significant correlations. Finally, we sorted motor neurons according to their weights in each source of inputs, such that motor neurons were assigned to the input with the highest relative strength. The accuracy of the classification was estimated as the percentage of motor neurons correctly assigned to the source of correlated inputs with the highest weight in each simulation.

### Application on experimental data

We finally representatively demonstrated the use of this approach for motor neuron spike trains experimentally recorded in humans and published in previous papers (14, 36). The detailed description of the experimental tasks and the procedures to collect and analyse data is available in each of these publications along with the datasets in public repositories. The first application consisted of a submaximal isometric plantarflexion while high-density electromyographic signals were recorded over the gastrocnemius lateralis and gastrocnemius medialis muscles (36). The second application consisted of a submaximal isometric dorsiflexion while high-density electromyographic signals were recorded over the tibialis anterior muscle. The third application consisted of a submaximal isometric knee extension while high-density electromyographic signals were recorded over the vastus lateralis and vastus medialis muscles (36). The fourth and last application consisted of an isometric multi-joint task where a participant pushed over a fixed pedal while high-density electromyographic signals were recorded over the gastrocnemius lateralis, gastrocnemius medialis, biceps femoris, semitendinosus, vastus lateralis, and vastus medialis (14). In all of these experiments, the signals were recorded using adhesive grids of 64 electrodes (13 × 5; gold coated; 1 mm diameter; 8 mm IED; OT Bioelettronica, Italy) placed over the muscle bellies. The skin was shaved and cleansed with an abrasive pad. Electrode-to-skin contact was maintained with a biadhesive perforated foam layer filled with conductive paste. Signals were recorded in monopolar mode with a sampling frequency of 2,048 Hz, amplified (x150), band-pass filtered (10–500 Hz), and digitised using a 400 channels acquisition system with a 16-bit resolution (EMG-Quattrocento; OT Bioelettronica, Italy).

Once recorded, the data were decomposed using convolutive blind-source separation (37). In short, a fixed-point algorithm that maximized the sparsity was applied to identify the sources embedded in the electromyographic signals, i.e., the motor unit spike trains. Motor unit spike trains can be considered as sparse sources with most samples being 0 (i.e., absence of spikes) and a few samples being 1 (i.e., spikes). In this algorithm, a contrast function was iteratively applied to the EMG signals to estimate the level of sparsity of the identified source, and the convergence was reached once the level of sparsity did not vary when compared to the previous iteration, with a tolerance fixed at 10^−4^ [see (37) for the definition of the detailed contrast functions]. At this stage, the estimated source contained high peaks (i.e., the spikes from the identified motor unit) and low peaks from other motor units and noise. High peaks were separated from low peaks and noise using peak detection and K-mean classification with two classes. The peaks from the class with the highest centroid were considered as the spikes of the identified motor unit. This decomposition procedure has been previously validated using experimental and simulated signals (37). After the automatic identification of the motor units, duplicates were removed, and all the motor unit spike trains were visually checked for false positives and false negatives (38). We ultimately applied the approach introduced in this paper to the firing activity of the identified motor units to representatively show the resulting graphs and clusters.

## Results

### Estimation of inputs to motor neurons

Our preliminary analyses aimed at determining whether the significant correlations between smoothed discharge rates accurately reflected the presence of correlated input between motor neurons. To this end, we used the simulations with N = 2 inputs. We report these results for durations ranging from 10s to 60s, as trial durations can vary during experiments and potentially impact the correlation values. We first calculated the significance thresholds defined as the 99^th^ percentile of the cross-correlation coefficient distribution generated with resampled versions of all motor neuron spike trains. These thresholds were set at 0.37, 0.26, 0.21, 0.18, 0.17, and 0.15 for 10s, 20s, 30s, 40s, 50s, and 60s, respectively. Figure 3A depicts the distribution of motor neuron pairs with significant and non-significant correlations as a function of their difference in weight of input #1. On average, pairs of motor neurons which did not exhibit correlated activity had a large difference in weights of 0.87 ± 0.13, 0.89 ± 0.11, 0.88 ± 0.12, 0.90 ± 0.10, 0.91 ± 0.09, and 0.92 ± 0.09 for 10s, 20s, 30s, 40s, 50s, and 60s, respectively; 1 being the maximal difference. This means that they received a small proportion of common inputs. Importantly, motor neurons that received only uncorrelated inputs, i.e., with a weight difference equal to 1, never exhibited significant correlation between their discharge rates, as expected theoretically. Moreover, we observed that the trial duration barely impacted the discrimination of pairs of motor neurons receiving common inputs, i.e., 91,9% of the pairs of motor neurons were either always or never correlated across the six durations. Consequently, the shape of force-directed graphs constructed using the matrix of significant correlations was similar across durations (Figure 3B). We performed all the following analyses using a trial duration of 60s.

**Figure 3:**
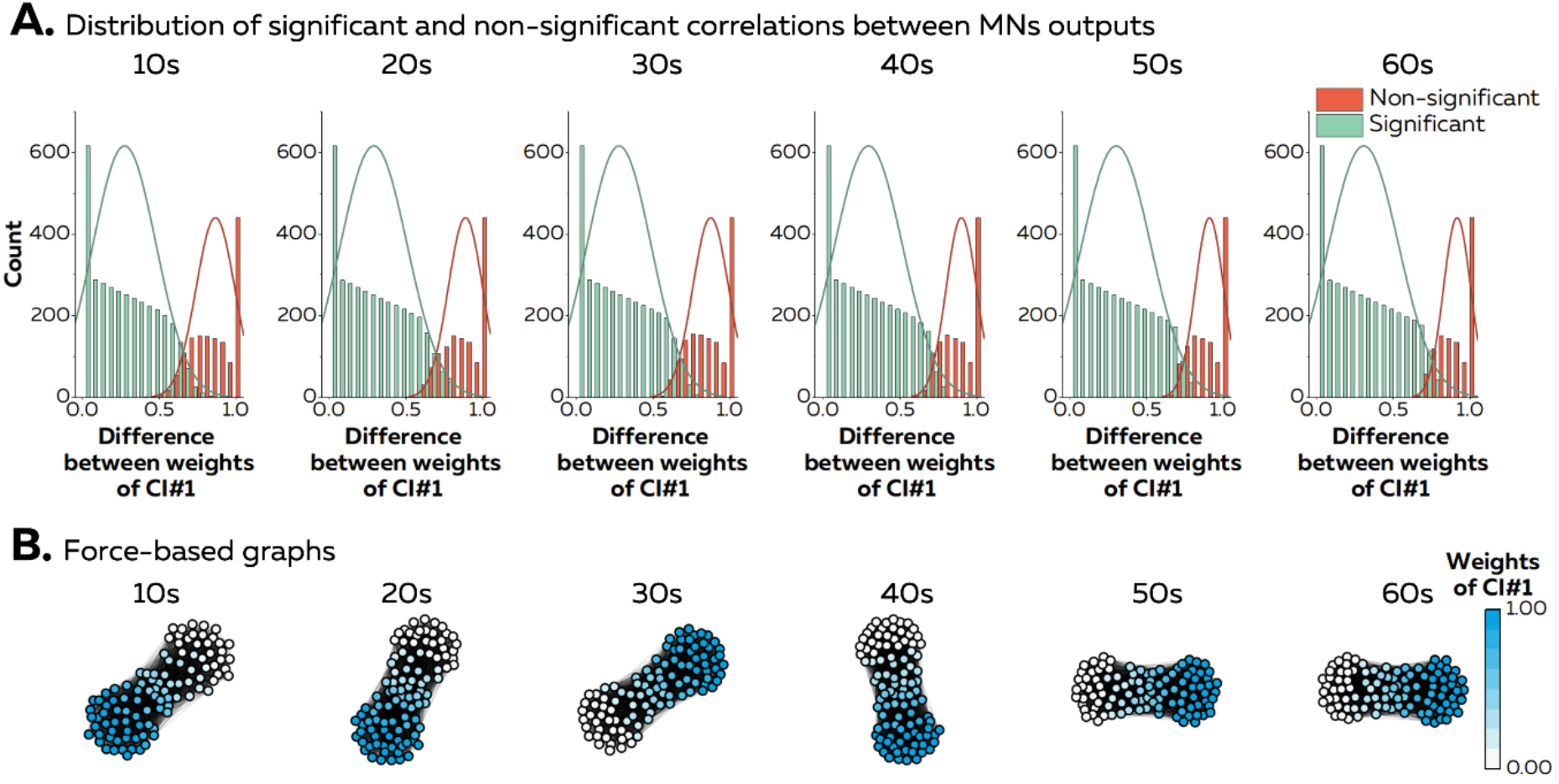
Accuracy of the estimation of correlated inputs from correlated smoothed discharge rates. (A) We reported the distribution of differences in weights of input 1 for all the pairs of motor neurons with (green) and without (red) significant correlations between their smoothed discharge rates. The lines represents a Gaussian distribution fitted to the data distribution. This analysis was repeated with correlations computed on segments of 10 to 60s (panels from left to right). (B). Force-directed graphs generated from the matrix of correlations calculated over segments of 10 to 60s. The color scheme depends on the relative strength of the input 1 for each motor neuron.

### Estimation of the number of inputs

We generated force-directed graphs using the algorithm developed by Fruchterman and Reingold (30). One property of this algorithm is that nodes, i.e., motor neurons, are positioned evenly in a two-dimensional space. In other words, groups of motor neurons loosely connected due to the presence of minimal, or no, correlated inputs tend to be pushed toward the edges of the space while keeping an equal distance between them. Thus, the shape of the graph gives a sense of the number of inputs received by the pool of motor neurons (Figure 4A). We applied a hierarchical clustering method to identify these groups of motor neurons loosely connected, and to estimate the number of inputs (Figure 4B). We identified 2, 3, and 4 clusters over the population of 100 motor neurons for simulation of 2, 3, and 4 inputs. In addition, when applied to the simulation where motor neurons received fully independent inputs, the force-directed graph approach did not identify any clear structure, with 11 clusters, and 17 motor neurons not connected to the graph (Figure 4A and B).

**Figure 4:**
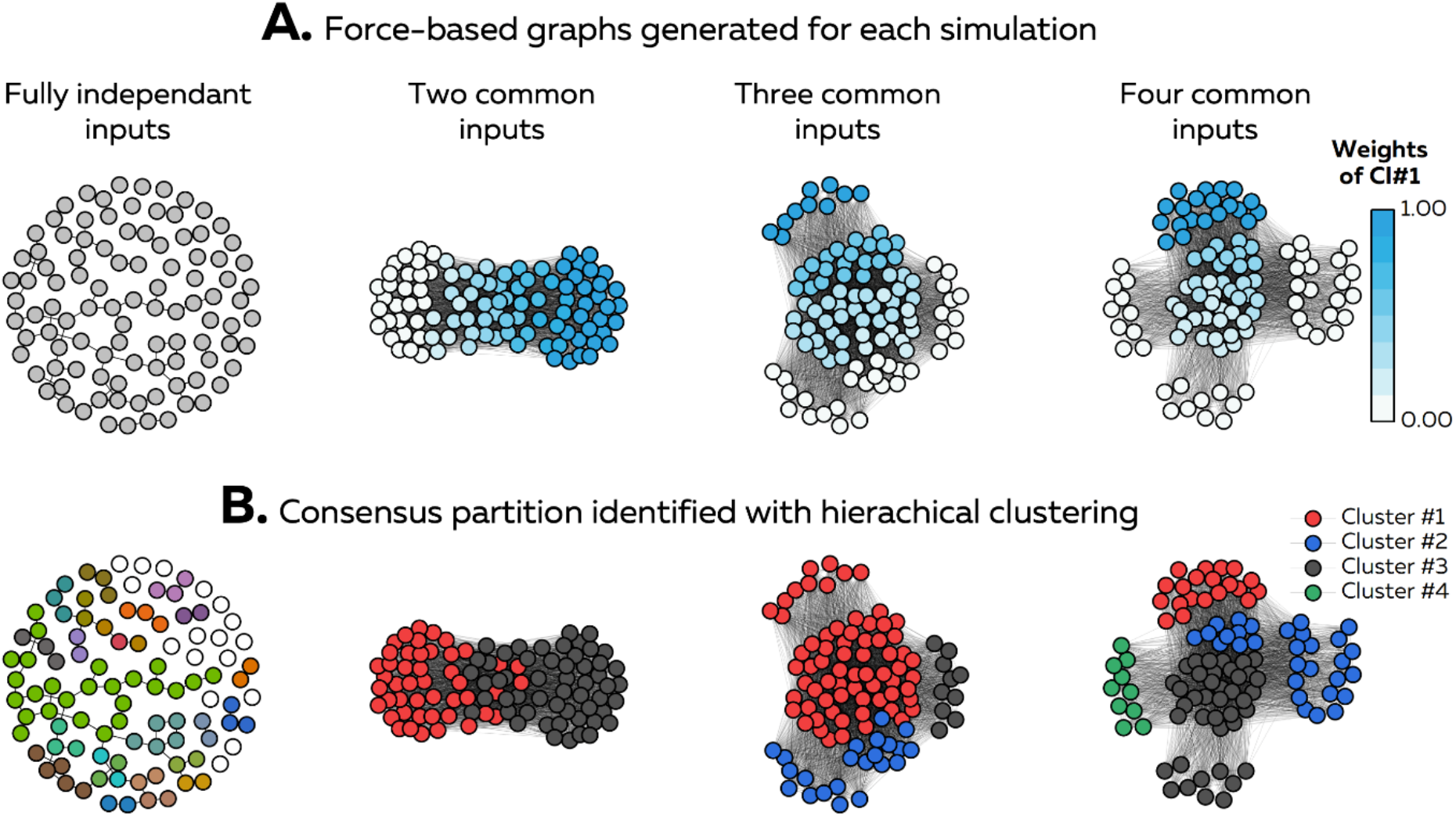
Force directed graphs and hierarchical clustering. (A) Force-directed graphs were generated from the adjacency matrices using the algorithm of Fruchterman and Reingold (30). The shapes of the graphs depended on the number of simulated inputs. The color scheme depended on the relative strength of input 1 for each motor neuron. (B) A hierarchical clustering algorithm was used to group motor neurons densely connected to each other and separate groups of motor neurons loosely connected (32). The color of each motor neuron depends on the cluster it belongs to.

After clustering the motor neurons, we used an iterative algorithm that identifies groups of motor neurons fully independent from the other clusters (Figure 5A; condition “0%“). On average, these refined clusters grouped motor neurons with the relative weight of one the sources of common input reaching 0.98 ± 0.04, 1.0 ± 0.0, and 0.99 ± 0.05 for graphs with 2, 3, and 4 sources of inputs, respectively. Given that a value of 1 means that the motor neuron received only a single source of correlated inputs, each cluster grouped motor neurons sharing almost 100% of the same source of common input. We then assessed the robustness of this method to estimate the number of inputs by perturbating motor neurons spike trains. Specifically, we removed 1% to 9% of the total number of firing instances or added false firings instances equal to 1% to 9% of the initial number of firing instances per motor neuron. The number of inputs was accurately identified for all the perturbed conditions for 2 inputs, for all the perturbed conditions except -8% and +8% for 3 inputs, and from -6% to +9% for 4 inputs (Figure 5B).

**Figure 5:**
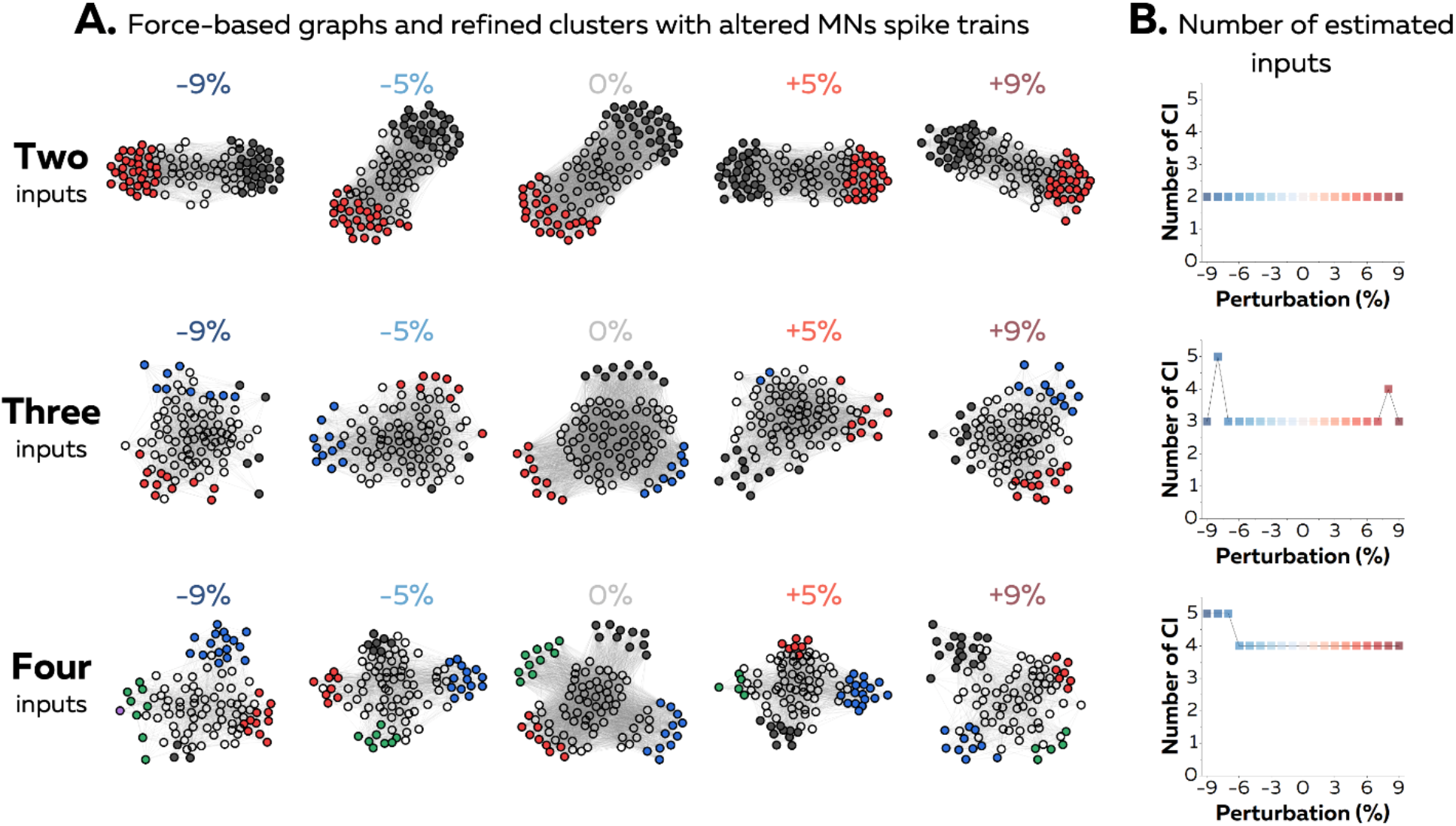
Robustness of the graph based-approach. We generated noisy versions of the motor neuron spike trains by randomly adding 1% to 9% of the intial number of spikes (false positives) or by randomly removing 1% to 9% of the initial number of spikes (fasle negatives). These errors mimic the typical error of the decomposition of electromyographic signals. (A) Force-directed graphs were generated from the correlation between these noisy version of the original spikes trains, and the graph-based approach was applied to estimate the number of inputs (B). Each color represents a cluster. The white nodes represent the motor neurons removed during our iterative procedure to identify groups of motor neurons fully independent from the other clusters.

### Estimation of the relative strengths of the inputs and motor neuron classification

For this analysis we only considered the conditions where the right number of inputs was estimated for all the simulations, i.e., from -5% to +5%. The classification of each motor neuron according to the cluster with the highest weight was barely affected by the perturbation of motor neuron spike trains for the three simulation. The accuracies were 99.2 ± 0.6%, 94.3 ± 2.2 %, and 95.1 ± 2.7 %, for 2, 3, and 4 inputs, respectively (Figure 6C). When considering the condition without any perturbation of the spike trains (0%), the RMSE of the estimated weights reached 0.11, 0.18, and 0.28 for 2, 3, and 4 inputs, respectively (Figure 6B). With 2 inputs, RMSE values changed by 0.01 ± 0.01 when we removed up to 5% of the discharge instances or added up to 5% of false discharge instances. With 3 inputs, the RMSE values slightly changed 0.02 ± 0.01. With 4 inputs, the RMSE changed by 0.02 ± 0.01. Overall, these results demonstrate the robustness of the graph-based approach to perturbations of the spike trains.

**Figure 6:**
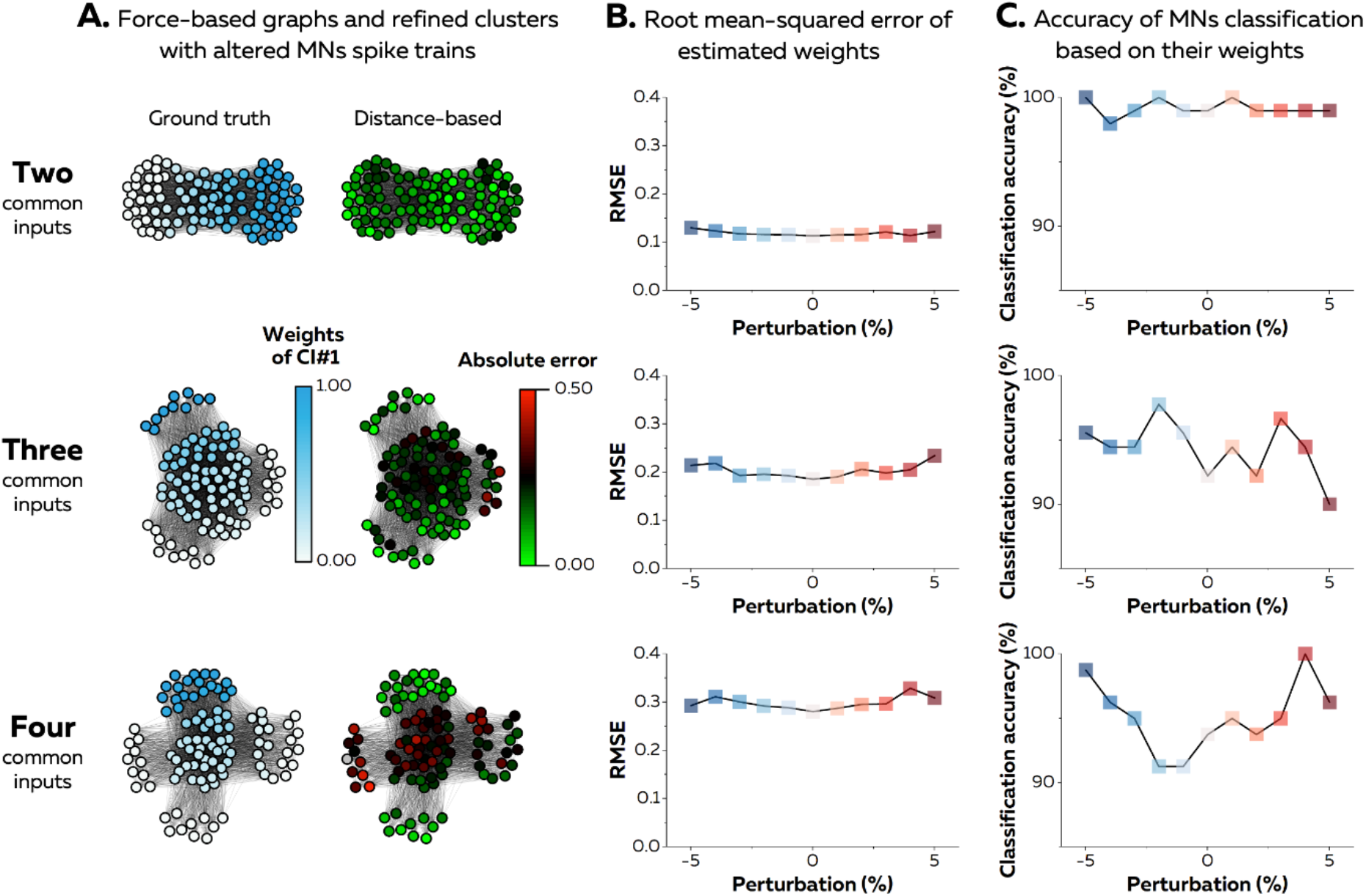
Estimation of the relative strength of the inputs for each motor neuron. We used the position of each motor neuron within the graph to estimate the relative strength of each input they receive. After identifying goups of motor neurons fully independent from the other clusters, we calculated their centroid. We considered the relative strength of each input as the distance separating each motor neuron from these centroids. We normalized the distances between 0 and 1 and subtracted them from 1, given that the shorter the distance, the higher the number of significant correlations. (A) Examples of the weighting of the input 1 for simulations with 2, 3, and 4 inputs. On the distance based panel, the color scheme representes the absolute error for each motor neuron. (B) Root mean squared error (RMSE) of the estimated weights for the graphs generated from the original adjacency matrix (0%), and for the graphs generated from spike trains contaminated by adding 1 to 5% of the original number of spikes (1 to 5) or by removing 1 to 5% of the original number of spikes (−1 to -5). (C). Classification of each motor neurons with the input having the highest relative strength. For example, if a given motor neuron has weights equal to 0.7, 0.1, 0.1, and 0.1 for inputs 1, 2, 3, and 4, respectively, it must me classified with the input 1.

### Impact of the proportion of identified motor neurons

We tested the impact of the proportion of identified motor neurons on the estimation of the number of inputs and their relative weights (Figure 7). To this end, we randomly selected motor neurons out of the initial group of 100 motor neurons by steps of 10 motor neurons and repeated this procedure 50 times per size of graph (Figure 7A). For the sake of clarity, we only present the results for the simulation with N = 2 common inputs. Our approach always identified two inputs, independently of the number of motor neurons considered in the analysis. RMSE reached 0.11 ± 0.0. Decreasing the number of motor neurons barely impacted the RMSE, with values of 0.10 ± 0.01 and 0.11 ± 0.0 for graphs of 10 and 100 motor neurons, respectively (Figure 7B). When considering the classification of motor neurons into a cluster, an accuracy of 97.4 ± 2.4% was observed across conditions. Specifically, the accuracy ranged from 96.8 ± 5.2 % to 97.4 ± 5.4 % for graphs of 10 and 100 motor neurons, respectively (Figure 7C).

**Figure 7:**
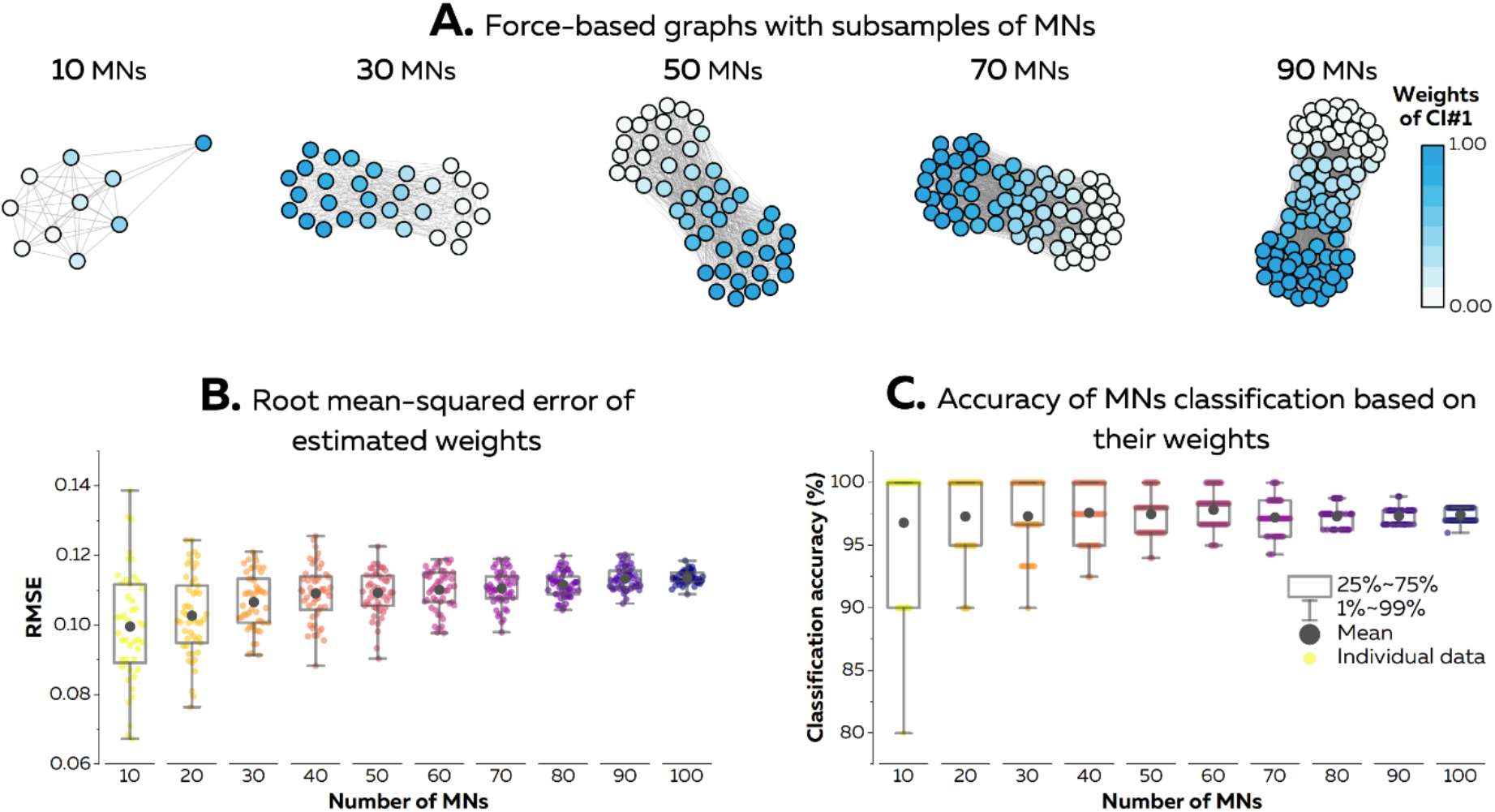
Impact of the number of identified motor neurons on the description of synergies. We randomly selected subsamples of motor neurons to identify the number of inputs and their relative strength from graphs of 10 to 100 motor neurons (A). The color scheme represents the relative strength of input 1 for each motor neuron. (B) Root mean squared error (RMSE) of the estimated weights using our approach, and (C) accuracy of the classification of each motor neuron with the input having the highest relative strength. Each dot represents an iteration, with 50 iterations per size of graph. The box represents 25% to 75% percentiles and the error bars 1% to 99% percentiles. MNs, motor neurons.

### Application to experimental data

We finally applied our approach to experimental data previously published and obtained from various lower leg muscles and various isometric motor tasks (14, 36) (Figure 8). For each dataset, only motor neurons with continuous firing activity during the isometric contraction were considered for the analysis. We first computed the cross-correlation coefficient between the smoothed discharge rates for each pair of motor neurons, and thresholded the adjacency matrix with a significance threshold calculated for each dataset (matrices in Figure 8). We then generated the force-directed graphs, applied hierarchical clustering, and used our approach to identify groups of motor neurons fully independent from the other clusters, as a way to estimate the number of common inputs to motor neurons.

**Figure 8:**
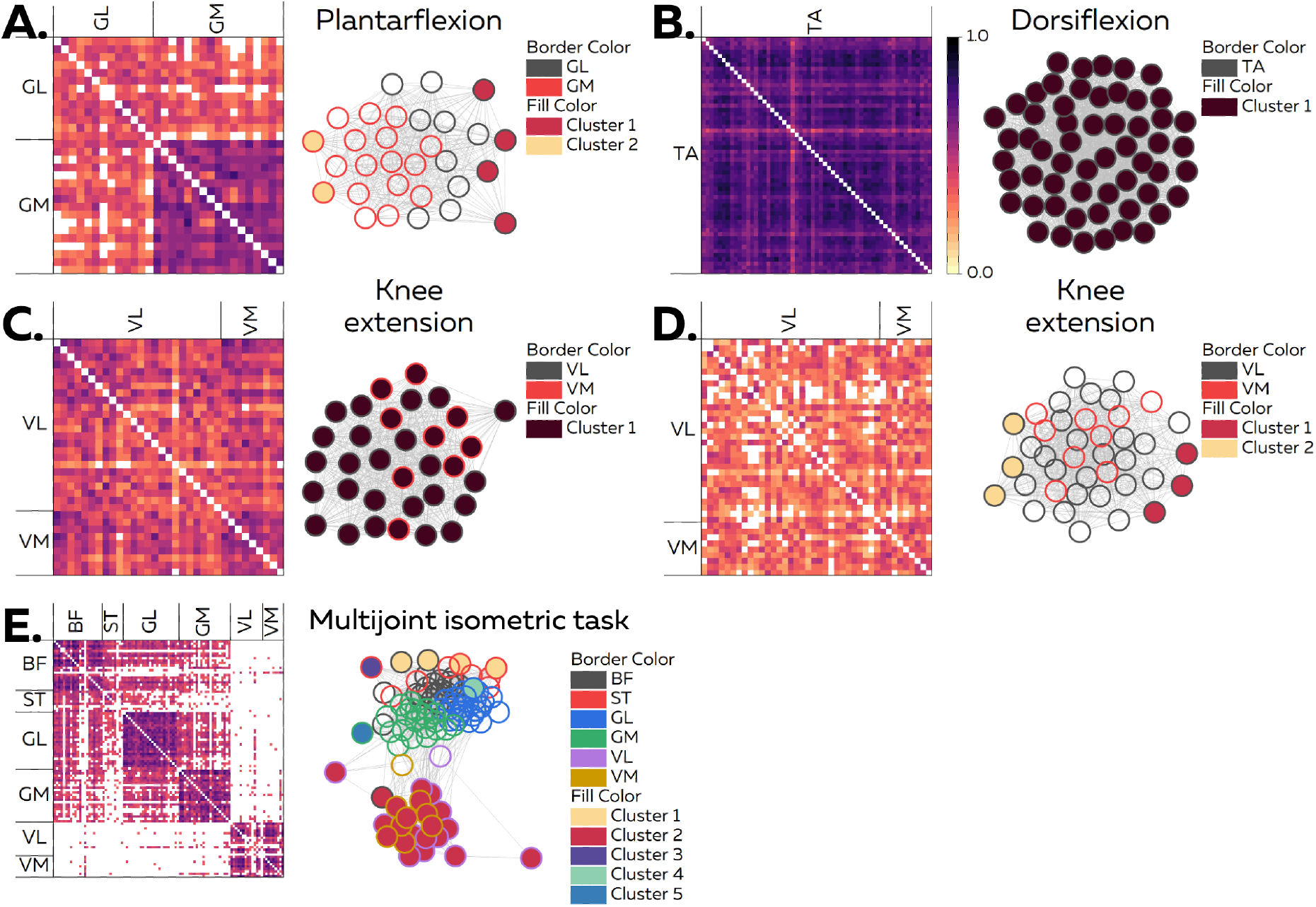
Application of the graph-based approach to experimental data. High-density electromyographic signals were recorded over the gastrocnemius lateralis (GL), gastrocnemius medialis (GM), tibialis anterior (TA), vastus lateralis (VL), vastus medialis (VM), biceps femoris (BF), and semitendinosus (ST) muscles. This data was decomposed using a blind source separation algorithm, and only motor neurons with a continuous firing activity during the isometric contraction were considered for the graph-based analysis. Each scatter is a motor neuron, and each line represents a significant correlation between two motor neurons. (A) We identified two clusters grouping motor neurons from the GL (Cluster 1) and the GM (Cluster 2). (B) Motor neurons from the TA were all grouped in the same cluster, with a high degree of correlation between all the motor neurons. (C) A unique cluster also grouped motor neurons from the VL and the VM for a participant. However, two clusters were identified for another participant (D). It is noteworthy than motor neurons innervating VL and VM were intermingled within the graph, and grouped in both clusters. (E) Finally, we identified five clusters grouping motor neurons from six muscles during an mutlijoint task. One cluster grouped motor neurons from VL and VM, while one cluster grouped motor neurons from the GL and one cluster grouped motor neurons from the GM. The two last clusters grouped motor neurons innervating both hamstring muscles, i.e., the BF and the ST.

For the ankle plantarflexion task, we identified two clusters of motor neurons innervating the gastrocnemius lateralis (GL) and gastrocnemius medialis (GM) muscles, with one cluster mostly grouping the motor neurons innervating the GL and the other cluster mostly grouping the motor neurons innervating the GM (Figure 8A). For the ankle dorsiflexion task, the smoothed discharge rates of all the motor neurons innervating the tibialis anterior were highly correlated, leading to the identification of only one cluster grouping all the identified motor neurons (Figure 8B). For the knee extension task, we observed that motor neurons innervating the vastus medialis and vastus lateralis muscles were intermingled within the graph (Figure 8C and 8D). In one participant, most of the motor neurons innervating the two muscles were highly correlated, leading to one cluster of motor neurons innervating the VL and VM muscles (Figure 8C). In a second participant, we observed less significant correlations between motor neurons, leading to two clusters, with motor neurons innervating both muscles in the two clusters (Figure 8D). Finally, we observed five clusters grouping motor neurons from six muscles in the lower limb during a multijoint isometric task (Figure 8E). Specifically, motor neurons innervating the VL and VM were grouped into the same cluster. Then, we observed one cluster gouping the motor neurons innervating the GL and one cluster gouping the motor neurons innervating the GM. Two other clusters were identified for the hamstring muscles (biceps femoris and semitendinosus muscles), each of these clusters grouping motor neurons innervating both muscles (Figure 8E).

## Discussion

We demonstrated the validity of a proposed graph-based approach to estimate the number of common inputs received by populations of motor neurons and their relative strength for each motor neuron, even with altered versions of the motor neuron spike trains. We also showcased how this approach could provide additional information on the neural control of motor neurons in humans. Together, these results open new perspectives to describe the number of dimensions controlled at the motor neuron level during and across human movements, helping researchers to support or oppose theories on the synergistic control of movements.

The foundation of this work relies on the concept of modularity of movement control (5, 6, 8), and the necessary transmission of common inputs to motor neurons to generate and modulate muscle forces (10, 25, 39, 40). We recently proposed that the central nervous system coordinates the firing activity of hundreds of spinal motor neurons during movements by transmitting a few correlated inputs to subgroups of these motor neurons (25). These ‘clusters’ can span over several muscles (14, 15, 23), and multiple common inputs can reach a single muscle or a part of it (14, 20, 21, 22, 23, 24). Based on these assumptions, here we presented a method that generates networks of correlated inputs to motor neurons by drawing graphs where nodes, i.e., motor neurons, are connected if they receive correlated inputs. To do so, we assessed the level of correlation between the smoothed discharge rates of each pair of motor neurons, and considered that they received correlated inputs if the correlation in their outputs was significant (14, 16, 29, 41). This approach largely attenuates the problem of the non-linearity between motor neuron inputs and outputs (29, 33, 42, 43, 44). Indeed, while the strength of correlation between inputs cannot be estimated from the correlation between outputs (42), a significant correlation between inputs very likely corresponds to a certain level of significant correlation between outputs. Thus, we limited the analysis of the output spike trains to the detection of the presence, not the strength, of common inputs. The validity of this assumption was directly tested by proving i) that pairs of motor neurons with a significant correlation always received most of the same inputs (Figure 3), and ii) that no clear graph structures emerged from motor neurons only receiving independent inputs (Figure 4). Interestingly, the duration of the signals used to compute the correlations did not impact the distribution of significant correlations across pairs of motor neurons nor the shape of the graphs (Figure 3).

We also found that our refinement of clusters, with the identification of groups of motor neurons that do not share any connections with the other clusters, can discriminate motor neurons that exclusively receive the same single source of common input (Figure 5, ‘Condition 0%). This is particurlary interesting to quantify the sources of common inputs within the population of identified motor neurons, i.e., the dimensionality of the neural control of motor neurons. Indeed, whether this ‘common drivers’ are consistent across human behaviors, or change depending on the mechanical contrains of the task, would determine their functional relevance to build human behaviors (1, 45). Additionally, a number of sources of common input significantly lower than the number of identified motor neurons would support the theory of synergies (8, 14, 22, 23). Conversely, having a number of ‘common drivers’ in the same order of dimension as the number of identified motor neurons woud support the hypothesis of a flexible control of single motor neurons (24, 46, 47).

Overall, our results made us confident that our approach can accurately map the distribution of correlated inputs across large datasets of motor neurons receiving several sources of common inputs. Moreover, the proposed graph-based method can also be interpreted as a way of visualizing high-dimensional spaces into two-dimensions. Indeed, conventional plots cannot display datasets with more than three dimensions. It comes as an alternative of several methods previously proposed in the field of computational neuroscience (48), biomechanics (49), or machine learning (50) to solve this issue, even if the output plots of the lattest methods can be abstract and difficult to interpret without complementary information.

The second step consisted in clustering motor neurons densely connected to each other while separating groups of motor neurons loosely connected. Many approaches exist to solve this problem (51), such as Isomap using the method of random walks (52), or the Louvain algorithm that optimizes the modularity (i.e., a metric that quantifies the strength of the clustering by comparing the number of connections within and between clusters (53)). They all have limitations due to stochastic processes or arbitrary choices of parameters (e.g., the resolution of the clusters; (54, 55)). This means that different clusters can emerge after each iteration, or that the optimization process stops at different sizes of clusters. One way to bypass these issues is to cluster the graph at multiple resolutions, going from one partition including all the motor neurons to smaller clusters (32, 56). The addition of consensus clustering, which identifies the partition with the clusters having the highest probability to be found across all the partitions generated by multiple iterations or multiple resolutions, also improves the robustness of the clustering (32, 34). Therefore, we used this method to accurately identify the number of simulated correlated inputs to motor neurons (Figures 4 and 5). Once refined, these clusters only grouped motor neurons that received a single source of the correlated inputs (Figure 5). The application of the proposed clustering approach to representative experimental data highlighted consistent results with previous studies that used coherence analyses or factorization algorithms. For example, we reported either one common input to both VL and VM muscles (11, 13), or two common inputs to intermingled motor neurons innervating each muscle (23). However, the scruture of the networks can sometime be noisier with a large number of clusters grouping a few motor neurons, as observed for some participants (see Supplementary Figure 1). Complementarity metrics that caracterize clusters as ‘strong’ or ‘weak’, such as the conductance (57), may help researchers to remove the weakest clusters, and thus select the right level of resolution within the multiple levels detected by hierarchical clustering (32).

Besides the estimation of the number of inputs, we also proposed a distance-based metric to estimate the relative strength of these inputs for each motor neuron (Figures 6 and 7). Dimensionality reduction on large scale neural recordings is usually achieved using approaches such as principal component analysis (22, 23), non-negative matrix factorization (58), or factorization algorithms (23). These algorithms usually identify neural latents, with time courses and weights, that can explain most of the variance of the recorded motor neuron spiking activity (59). One caveat of such approaches is the necessity of identifying the number of neural latents a priori, using an arbitrary threshold on the level of variance explained by the model (7, 59). Alternatively, a lengthy and computationally demanding approach consists of identifying the number of latents that maximizes the cross-validated data likelihood with probabilistic methods (60). Conversely, the proposed approach first estimates the number of inputs and then uses a simple metric based on the Euclidean distance between each node and the centroid of each refined cluster to estimate the relative strength of each input to each motor neuron (Figures 6 and 7). Importantly, our method was robust to the perturbation of the spiking activity of individual motor neurons that mimic the errors of automatic electromyographic decomposition (35, 38). Additonally, the root-mean squared error in the estimate of input gains did not vary when decreasing the number of motor neurons used to build the graph (Figure 7). This is particularly important when considering that currently it is only possible to experimentally identify a subset of the active motor neurons during natural tasks. As such, this method could add a layer of information for reasearchers trying to quantify and describe the dimensionality of the neural control of motor neurons during and across human movements.

## References

1. d’Avella A, Saltiel P, Bizzi E. Combinations of muscle synergies in the construction of a natural motor behavior. Nat Neurosci. 2003;6(3):300–8.

2. Cheung VC, d’Avella A, Tresch MC, Bizzi E. Central and sensory contributions to the activation and organization of muscle synergies during natural motor behaviors. J Neurosci. 2005;25(27):6419–34.

3. Dominici N, Ivanenko YP, Cappellini G, d’Avella A, Mondi V, Cicchese M, et al. Locomotor primitives in newborn babies and their development. Science. 2011;334(6058):997–9.

4. Takei T, Confais J, Tomatsu S, Oya T, Seki K. Neural basis for hand muscle synergies in the primate spinal cord. Proc Natl Acad Sci U S A. 2017;114(32):8643–8.

5. Tresch MC, Jarc A. The case for and against muscle synergies. Curr Opin Neurobiol. 2009;19(6):601–7.

6. Ting LH, Chiel HJ, Trumbower RD, Allen JL, McKay JL, Hackney ME, et al. Neuromechanical principles underlying movement modularity and their implications for rehabilitation. Neuron. 2015;86(1):38–54.

7. Tresch MC, Cheung VC, d’Avella A. Matrix factorization algorithms for the identification of muscle synergies: evaluation on simulated and experimental data sets. J Neurophysiol. 2006;95(4):2199–212.

8. Cheung VCK, Seki K. Approaches to revealing the neural basis of muscle synergies: a review and a critique. J Neurophysiol. 2021;125(5):1580–97.

9. Overduin SA, d’Avella A, Carmena JM, Bizzi E. Microstimulation activates a handful of muscle synergies. Neuron. 2012;76(6):1071–7.

10. Farina D, Negro F, Muceli S, Enoka RM. Principles of Motor Unit Physiology Evolve With Advances in Technology. Physiology (Bethesda). 2016;31(2):83–94.

11. Laine CM, Martinez-Valdes E, Falla D, Mayer F, Farina D. Motor Neuron Pools of Synergistic Thigh Muscles Share Most of Their Synaptic Input. J Neurosci. 2015;35(35):12207–16.

12. Del Vecchio A, Falla D, Felici F, Farina D. The relative strength of common synaptic input to motor neurons is not a determinant of the maximal rate of force development in humans. J Appl Physiol (1985). 2019;127(1):205–14.

13. Avrillon S, Del Vecchio A, Farina D, Pons JL, Vogel C, Umehara J, et al. Individual differences in the neural strategies to control the lateral and medial head of the quadriceps during a mechanically constrained task. J Appl Physiol (1985). 2021;130(1):269–81.

14. Hug F, Avrillon S, Sarcher A, Del Vecchio A, Farina D. Correlation networks of spinal motor neurons that innervate lower limb muscles during a multi-joint isometric task. J Physiol. 2022.

15. Gibbs J, Harrison LM, Stephens JA. Organization of inputs to motoneurone pools in man. J Physiol. 1995;485 (Pt 1)(Pt 1):245–56.

16. De Luca CJ, Erim Z. Common drive in motor units of a synergistic muscle pair. J Neurophysiol. 2002;87(4):2200–4.

17. Negro F, Holobar A, Farina D. Fluctuations in isometric muscle force can be described by one linear projection of low-frequency components of motor unit discharge rates. J Physiol. 2009;587(Pt 24):5925–38.

18. Negro F, Yavuz US, Farina D. The human motor neuron pools receive a dominant slow-varying common synaptic input. J Physiol. 2016;594(19):5491–505.

19. Muceli S, Poppendieck W, Holobar A, Gandevia S, Liebetanz D, Farina D. Blind identification of the spinal cord output in humans with high-density electrode arrays implanted in muscles. Science advances. 2022;8(46):eabo5040.

20. Keen DA, Fuglevand AJ. Common input to motor neurons innervating the same and different compartments of the human extensor digitorum muscle. J Neurophysiol. 2004;91(1):57–62.

21. McIsaac TL, Fuglevand AJ. Motor-unit synchrony within and across compartments of the human flexor digitorum superficialis. J Neurophysiol. 2007;97(1):550–6.

22. Madarshahian S, Letizi J, Latash ML. Synergic control of a single muscle: The example of flexor digitorum superficialis. J Physiol. 2021;599(4):1261–79.

23. Del Vecchio A, Germer C, Kinfe TM, Nuccio S, Hug F, Eskofier B, et al. Common synaptic inputs are not distributed homogeneously among the motor neurons that innervate synergistic muscles. bioRxiv. 2022:2022.01.23.477379.

24. Marshall NJ, Glaser JI, Trautmann EM, Amematsro EA, Perkins SM, Shadlen MN, et al. Flexible neural control of motor units. Nat Neurosci. 2022;25(11):1492–504.

25. Hug F, Avrillon S, Ibanez J, Farina D. Common synaptic input, synergies and size principle: Control of spinal motor neurons for movement generation. J Physiol. 2023;601(1):11–20.

26. Boonstra TW, Danna-Dos-Santos A, Xie HB, Roerdink M, Stins JF, Breakspear M. Muscle networks: Connectivity analysis of EMG activity during postural control. Sci Rep. 2015;5:17830.

27. Kerkman JN, Daffertshofer A, Gollo LL, Breakspear M, Boonstra TW. Network structure of the human musculoskeletal system shapes neural interactions on multiple time scales. Science advances. 2018;4(6):eaat0497.

28. Elias LA, Kohn AF. Individual and collective properties of computationally efficient motoneuron models of types S and F with active dendrites. Neurocomputing. 2013;99:521–33.

29. Rodriguez-Falces J, Negro F, Farina D. Correlation between discharge timings of pairs of motor units reveals the presence but not the proportion of common synaptic input to motor neurons. J Neurophysiol. 2017;117(4):1749–60.

30. Fruchterman TMJ, Reingold EM. Graph Drawing by Force-Directed Placement. Software Pract Exper. 1991;21(11):1129–64.

31. Binder MD, Powers RK, Heckman CJ. Nonlinear Input-Output Functions of Motoneurons. Physiology (Bethesda). 2020;35(1):31–9.

32. Jeub LGS, Sporns O, Fortunato S. Multiresolution Consensus Clustering in Networks. Sci Rep. 2018;8(1):3259.

33. Farina D, Negro F, Dideriksen JL. The effective neural drive to muscles is the common synaptic input to motor neurons. J Physiol. 2014;592(16):3427–41.

34. Lancichinetti A, Fortunato S. Consensus clustering in complex networks. Sci Rep. 2012;2:336.

35. Holobar A, Minetto MA, Farina D. Accurate identification of motor unit discharge patterns from high-density surface EMG and validation with a novel signal-based performance metric. J Neural Eng. 2014;11(1):016008.

36. Rossato J, Tucker K, Avrillon S, Lacourpaille L, Holobar A, Hug F. Less common synaptic input between muscles from the same group allows for more flexible coordination strategies during a fatiguing task. J Neurophysiol. 2022;127(2):421–33.

37. Negro F, Muceli S, Castronovo AM, Holobar A, Farina D. Multi-channel intramuscular and surface EMG decomposition by convolutive blind source separation. J Neural Eng. 2016;13(2):026027.

38. Hug F, Avrillon S, Del Vecchio A, Casolo A, Ibanez J, Nuccio S, et al. Analysis of motor unit spike trains estimated from high-density surface electromyography is highly reliable across operators. J Electromyogr Kinesiol. 2021;58:102548.

39. De Luca CJ, Erim Z. Common drive of motor units in regulation of muscle force. Trends in neurosciences. 1994;17(7):299–305.

40. Farina D, Negro F. Common synaptic input to motor neurons, motor unit synchronization, and force control. Exerc Sport Sci Rev. 2015;43(1):23–33.

41. Maillet J, Avrillon S, Nordez A, Rossi J, Hug F. Handedness is associated with less common input to spinal motor neurons innervating different hand muscles. J Neurophysiol. 2022;128(4):778–89.

42. de la Rocha J, Doiron B, Shea-Brown E, Josic K, Reyes A. Correlation between neural spike trains increases with firing rate. Nature. 2007;448(7155):802–6.

43. Negro F, Farina D. Linear transmission of cortical oscillations to the neural drive to muscles is mediated by common projections to populations of motoneurons in humans. J Physiol. 2011;589(Pt 3):629–37.

44. Negro F, Farina D. Factors influencing the estimates of correlation between motor unit activities in humans. PLoS One. 2012;7(9):e44894.

45. d’Avella A, Bizzi E. Shared and specific muscle synergies in natural motor behaviors. Proc Natl Acad Sci U S A. 2005;102(8):3076–81.

46. Formento E, Botros P, Carmena JM. A non-invasive brain-machine interface via independent control of individual motor units. bioRxiv. 2021:2021.03.22.436518.

47. Basmajian JV. Control and training of individual motor units. Science. 1963;141(3579):440–1.

48. Cowley BR, Kaufman MT, Butler ZS, Churchland MM, Ryu SI, Shenoy KV, et al. DataHigh: graphical user interface for visualizing and interacting with high-dimensional neural activity. J Neural Eng. 2013;10(6):066012.

49. Valero-Cuevas FJ, Cohn BA, Yngvason HF, Lawrence EL. Exploring the high-dimensional structure of muscle redundancy via subject-specific and generic musculoskeletal models. J Biomech. 2015;48(11):2887–96.

50. Van der Maaten L, Hinton G. Visualizing data using t-SNE. Journal of machine learning research. 2008;9(11).

51. Fortunato S, Hric D. Community detection in networks: A user guide. Physics Reports-Review Section of Physics Letters. 2016;659:1–44.

52. Rosvall M, Bergstrom CT. Maps of random walks on complex networks reveal community structure. Proc Natl Acad Sci U S A. 2008;105(4):1118–23.

53. Blondel VD, Guillaume JL, Lambiotte R, Lefebvre E. Fast unfolding of communities in large networks. Journal of Statistical Mechanics-Theory and Experiment. 2008;2008(10):P10008.

54. Lancichinetti A, Fortunato S. Limits of modularity maximization in community detection. Phys Rev E Stat Nonlin Soft Matter Phys. 2011;84(6 Pt 2):066122.

55. Traag VA, Krings G, Van Dooren P. Significant scales in community structure. Sci Rep. 2013;3(1):2930.

56. Schaub MT, Delvenne JC, Yaliraki SN, Barahona M. Markov dynamics as a zooming lens for multiscale community detection: non clique-like communities and the field-of-view limit. PLoS One. 2012;7(2):e32210.

57. Leskovec J, Lang KJ, Dasgupta A, Mahoney MW. Community Structure in Large Networks: Natural Cluster Sizes and the Absence of Large Well-Defined Clusters. Internet Mathematics. 2009;6(1):29–123.

58. Tanzarella S, Muceli S, Santello M, Farina D. Synergistic Organization of Neural Inputs from Spinal Motor Neurons to Extrinsic and Intrinsic Hand Muscles. J Neurosci. 2021;41(32):6878–91.

59. Cunningham JP, Yu BM. Dimensionality reduction for large-scale neural recordings. Nat Neurosci. 2014;17(11):1500–9.

60. Yu BM, Cunningham JP, Santhanam G, Ryu SI, Shenoy KV, Sahani M. Gaussian-process factor analysis for low-dimensional single-trial analysis of neural population activity. J Neurophysiol. 2009;102(1):614–35.

